# NUDT18 catalyzes the hydrolysis of active metabolites of the antivirals Remdesivir, Ribavirin and Molnupiravir

**DOI:** 10.1101/2021.11.16.468802

**Authors:** Ann-Sofie Jemth, Emma Rose Scaletti, Evert Homan, Pål Stenmark, Thomas Helleday, Maurice Michel

## Abstract

Remdesivir and Molnupiravir have gained considerable interest due to their activity against SARS-CoV-2. Cellular hydrolysis of their active triphosphate forms, Remdesivir-TP and Molnupiravir-TP, would decrease drug efficiency. We therefore tested Remdesivir-TP as a substrate against a panel of human hydrolases and found that NUDT18 catalyzes the hydrolysis of Remdesivir-TP. The k_cat_ value of NUDT18 for Remdesivir-TP was determined to 2.6 s^-1^ and the K_m_ value was 156 μM, suggesting that NUDT18 catalyzed hydrolysis of Remdesivir-TP occurs in cells. We demonstrate that the triphosphates of the antivirals Ribavirin and Molnupiravir are hydrolyzed by NUDT18, albeit with a lower efficiency compared to Remdesivir-TP. NUDT18 also hydrolyses the triphosphates of Sofosbuvir and Aciclovir although with significantly lower activity. These results suggest that NUDT18 can act as a cellular sanitizer of modified nucleotides and may influence the antiviral efficacy of Remdesivir, Molnupiravir and Ribavirin. NUDT18 is expressed in respiratory epithelial cells and may limit the antiviral efficacy of Remdesivir and Molnupiravir against SARS-CoV2 replication by decreasing the intracellular concentration of their active metabolites at their intended site of action.

## Introduction

The recent SARS-CoV-2 pandemic with its fast worldwide spread and high mortality rate led to major efforts to repurpose already approved antiviral drugs in order to quickly control the outbreak and thereby save lives. Remdesivir (GS-5734) was originally developed for the treatment of Ebola but is a broad-acting antiviral which has been found to reduce the replication of coronaviruses both in cell culture and animal models [1]. Remdesivir treatment was shown to decrease the severity of disease after MERS-CoV infection via decreasing virus replication and thereby reducing damage to the lungs when administered both before and after infection in a rhesus macaque model [1]. Recently, Remdesivir was tested in several clinical trials for treatment of COVID-19 caused by the coronavirus SARS-CoV-2. Remdesivir was shown to shorten the recovery time of hospitalized adults with COVID-19 as well as reducing mortality rate [2] and suggested to be most effective if administered early after infection [3]. Remdesivir was the first drug to be approved for treatment of severe COVID-19 in July 2020 by the European Medicine Agency and later in the fall by the U.S Food and Drug Administration (FDA) [2, 4–6]. Remdesivir is a prodrug in the form of a protected phosphate that upon cell entry (Figure 1, step 1) is metabolized by cellular enzymes to the corresponding monophosphate (GS-441524, Figure 1, step 2). The monophosphate is subsequently phosphorylated by cellular kinases to its active triphosphate form (Remdesivir-TP, aka GS-443902, Figure 1, step 3). The antiviral effect of Remdesivir-TP is mediated through its incorporation into the growing virus RNA chain by the viral RNA-dependent RNA polymerase (RdRp, Figure 1, step 4). Incorporation of the Remdesivir metabolite causes the polymerase to stall and RNA replication to be terminated (Figure 1, step 5), leading to a decrease in the production of viral RNA [7, 8].

**Figure 1.**
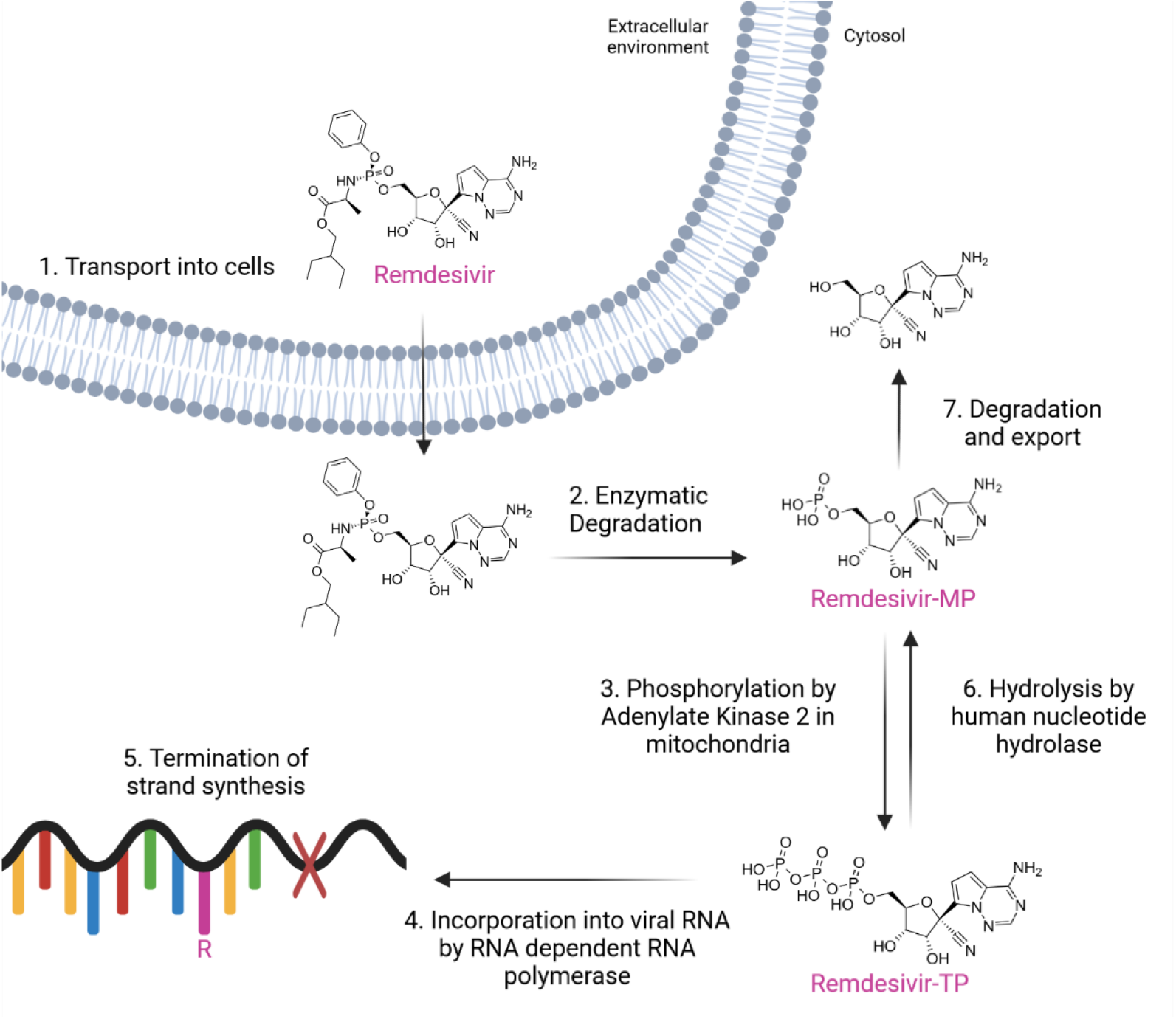
Structure, intracellular metabolism and mode of action of Remdesivir

Molnupiravir (EIDD-2801) was in November 2021 given a temporary authorization by the UK’s medicines regulatory authority for treatment of mild to moderate COVID-19 in adults at risk for severe disease and one month later an emergency use authorization was issued by the FDA. Administration of this orally active drug lead to a clear reduction of SARS-CoV-2 replication as well as quicker recovery time [9]. In addition, early treatment with Molnupiravir clearly reduced the risk of death or hospitalization in unvaccinated adults with COVID-19 at risk for severe disease [10]. Molnupiravir is a prodrug for β-d-N4-hydroxycytidine, which previously has been shown to be effective against various RNA viruses including influenza and other coronaviruses [11, 12], and is rapidly hydrolyzed to β-d-N4-hydroxycytidine followed by phosphorylation resulting in formation of its active 5’-triphosphate metabolite [13]. The incorporation of β-d-N4-hydroxycytidine triphosphate, hereafter called Molnupiravir-TP, by RdRp into viral RNA causes a drastic increase in the mutation frequency resulting in a decrease in SARS-CoV-2 replication [14].

Ribavirin is another antiviral drug that was tested for activity towards SARS-CoV-2. Ribavirin has since the 1970s been known to act as a broad-spectrum antiviral of both DNA and RNA viruses [15]. It has been used in combination with IFN-α-based therapy to treat hepatitis C [16] as well as other viral infections such as Lassa fever [17], Crimean–Congo fever [18, 19] and chronic hepatitis E virus infections [20]. Ribavirin in combination with IFN-β was shown to synergistically inhibit SARS-associated coronavirus replication in animal and human cell lines [21] suggesting that Ribavirin could be a potential treatment option for SARS-CoV-2. The precise mode of action of Ribavirin is not completely understood and its metabolites have been suggested to target several processes of which the antiviral impact may depend on their intracellular concentrations [22]. However, it is known that Ribavirin-MP inhibits inosine monophosphate dehydrogenase leading to GTP depletion with effects on many cellular processes. Ribavirin-TP has been hypothesized to directly inhibit viral RNA dependent polymerase and may at higher concentrations also cause mutations in the viral genome. Ribavirin-TP is hydrolyzed by the ITPase hydrolase and patients carrying alleles encoding low activity ITPase variants display a reduced risk of hepatitis C relapse, suggesting that these patients have higher concentrations of the active Ribavirin metabolite. Furthermore, low activity ITPase alleles have been shown to protect against anemia acquired upon INF-α and Ribavirin therapy [22, 23].

The ITPase activity-dependent outcome of Ribavirin treatment strongly emphasizes the impact of cellular metabolism on drug efficacy. Similar to ITPase, several NUDIX enzymes catalyze the hydrolysis of modified nucleoside triphosphates [24, 25] and have been shown to modulate the efficacy of various drugs including antivirals [26, 27]. An example is NUDT15 which we recently reported catalyzes the hydrolysis of Ganciclovir-TP [27]. In a similar way, hydrolysis of Remdesivir-TP into Remdesivir-MP, catalyzed by a human nucleotide hydrolase (Figure 1, Step 6) would reduce the cellular concentration of the active metabolite. Moreover, conversion of Remdesivir-MP to the corresponding nucleoside through the action of 5’-nucleotidase would enable transport out of the cell (Figure 1, Step 7) and further decrease the antiviral potency of Remdesivir. In order to investigate if such a hydrolase could be found within the NUDIX family of enzymes we tested Remdesivir-TP as substrate for a panel of human NUDIX hydrolases and found that NUDT18 catalyzed the hydrolysis of Remdesivir-TP. Encouraged by this we tested NUDT18 for activity towards several antiviral triphosphates and found NUDT18 to also be an efficient catalyst of Ribavirin-TP and Molnupiravir-TP hydrolysis. NUDT18 has to date only been described to display activity towards 8-oxo-(d)GDP and has therefore been suggested to cleanse the nucleotide pool from oxidatively damaged nucleotides, preventing their incorporation into DNA [28, 29]. The results of this study suggest that NUDT18 acts as a cellular sanitizer in a broader sense and contributes to removing modified nucleotides of both endogenous and exogenous origin, and potentially modulates the efficiency of the antiviral drugs Remdesivir, Molnupiravir and Ribavirin.

## Results and discussion

### NUDT18 catalyzes the hydrolysis of Remdesivir-TP and Ribavirin-TP

In order to investigate if human NUDIX enzymes can modulate the cellular concentration of the active metabolite of Remdesivir and thereby reduce its efficacy as antiviral drug we tested human MTH1, NUDT5, NUDT9, NUDT15 and NUDT18 for hydrolysis activity with Remdesivir-TP. Among these NUDIX hydrolases only NUDT18 was found to catalyze the hydrolysis of Remdesivir-TP (Figure 2A). NUDT18 was shown to catalyze the hydrolysis of both the bond between the α- and β-phosphate groups, generating PPi, as well as the bond between the β- and γ-phosphate groups, generating Pi (Figure 2A and B). This was evidenced by an increase in the amount of Pi formed in the presence of PPase, which hydrolyzes PPi. Based on the described activity of ITPase with Ribavirin-TP, which constitutes one of the active metabolites of the antiviral Ribavirin, we argued that ITPase may also hydrolyze other antiviral triphosphates. We therefore tested human ITPase as well as the human hydrolases dCTPase and dUTPase for activity with Remdesivir-TP. Human dCTPase displayed low but measurable activity with Remdesivir-TP while no activity was detected with dUTPase or ITPase (Figure 2C). Comparison with NUDT18 activity towards 8-oxo-dGDP showed an approximately 2-fold higher specific activity compared to Remdesivir-TP at 100 μM substrate concentration (Figure 2C).

**Figure 2.**
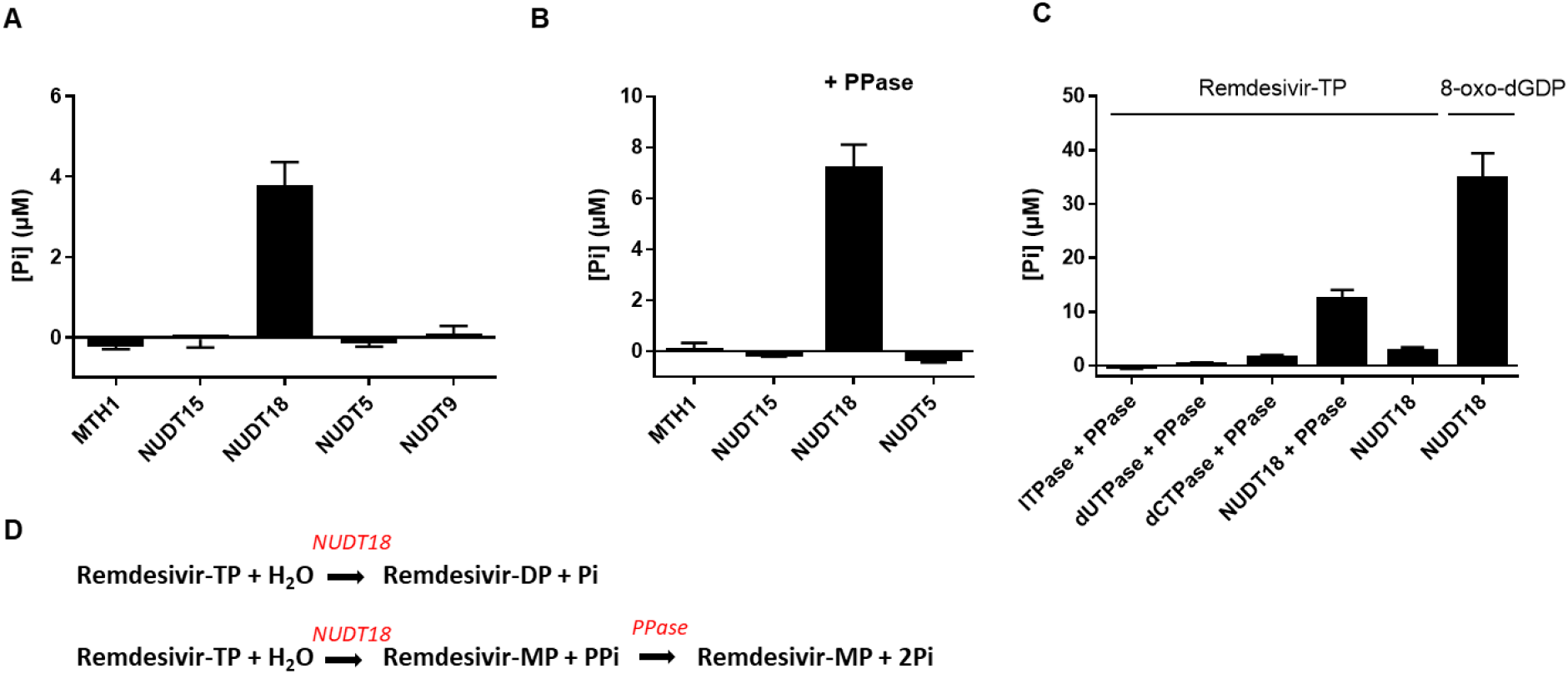
NUDT18 catalyzes Remdesivir-TP hydrolysis. **(A)** Hydrolysis activity of MTH1, NUDT15, NUDT18, NUDT5 and NUDT9 (200 nM) with 100 μM Remdesivir-TP was tested in assay buffer (20 mM TrisAcetate pH 8.0, 40 mM NaCl, 10 mM MgAc, 1 mM DTT, 0.05% Tween20) by incubation at 22 °C for 30 minutes. Formed Pi was detected by addition of malachite green reagent and measurement of absorbance at 630 nm. **(B)** Activity of 200 nM MTH1, NUDT15, NUDT18, NUDT5, was also tested in presence of PPase (0.2 U/ml) that converts formed PPi to Pi. **(C)** Activity of the human pyrophoshatases ITPase, dUTPase and dCTPase with 100 μM Remdesivir-TP was assayed in presence of PPase was assayed and compared to activity of NUDT18 with and without PPase and to NUDT18 activity with 100 μM 8-oxo-dGDP. **(D)** Hydrolysis products of NUDT18 catalyzed hydrolysis of Remdesivir-TP in presence and absence of NUDT18.

The observed activity of NUDT18 with Remdesivir-TP prompted us to test other antiviral triphosphates as substrates for NUDT18. We found that Ribavirin-TP is also hydrolyzed by NUDT18, albeit not to the same extent as Remdesvir-TP (Figure 3A). Similar to Remdesivir-TP, NUDT18 catalyzed Ribavirin-TP hydrolysis results in the production of both Ribavirin di- and monophosphate, as shown by an increase in the signal when PPase was included in the assay buffer (Figure 3A). NUDT18 was also shown to hydrolyze Molnupiravir-TP. As evidenced by a 5-fold higher production of Pi in the presence of PPase (Figure 3B), Molnupiravir-TP hydrolysis of the bond between the α- and β-phosphate groups appears to be preferred. In addition, NUDT15, previously shown to catalyze the hydrolysis of Ganciclovir-TP [27], was found to catalyze the hydrolysis of Molnupiravir-TP producing pyrophosphate, albeit with considerably lower efficiency compared to NUDT18, while MTH1 did not display any detectable activity with this substrate (Figure 3B). To investigate the activity of NUDT18 with these substrates in more detail we performed kinetic analyses of the NUDT18-catalyzed hydrolysis of Remdesivir-TP, Ribavirin-TP, Molnupiravir-TP and for comparison with 8-oxo-dGDP. Enzyme activities were assayed under identical conditions and enzyme saturation curves using these substrates were produced (Figure 4A, B, C and D) and kinetic parameters were determined (Figure 4G). The K_m_ value for NUDT18 was determined to be 16.5 μM for 8-oxo-dGDP, which is in agreement with a previously published K_m_ value of 12 μM [28]. The k_cat_ value for 8-oxo-dGDP at 22°C was determined to be 1.8 s^-1^ and the calculated k_cat_/K_m_ value was 109,800 M^-1^s^-1^. The K_m_ value for Remdesivir-TP was determined to be 156 μM (n=2) and the k_cat_ value was 2.6 s^-1^ resulting in a k_cat_/K_m_ value of 17,700 M^-1^s^-1^. The approximately 6-fold higher catalytic efficiency (k_cat_/K_m_ value) of NUDT18 for 8-oxo-dGDP means that 8-oxo-dGDP would be the preferred substrate over Remdesivir-TP at the same concentration. However, the concentration of Remdesivir in monkey lung cell tissue post IV administration was estimated to be 800 nM [30]. Based on an estimation of the intracellular 8-oxo-dGTP concentration of 2 nM in U2OS cells [31], and the assumption that the cellular concentration of 8-oxo-dGDP is likely in a similar range or lower, it is likely that the cellular Remdesivir-TP concentration is at least 100-fold higher than the concentration of 8-oxo-dGDP *in vivo*. This suggests that the activity of NUDT18 with Remdesivir-TP may be of relevance for modulating the Remdesivir-TP concentration in cells.

**Figure 3.**
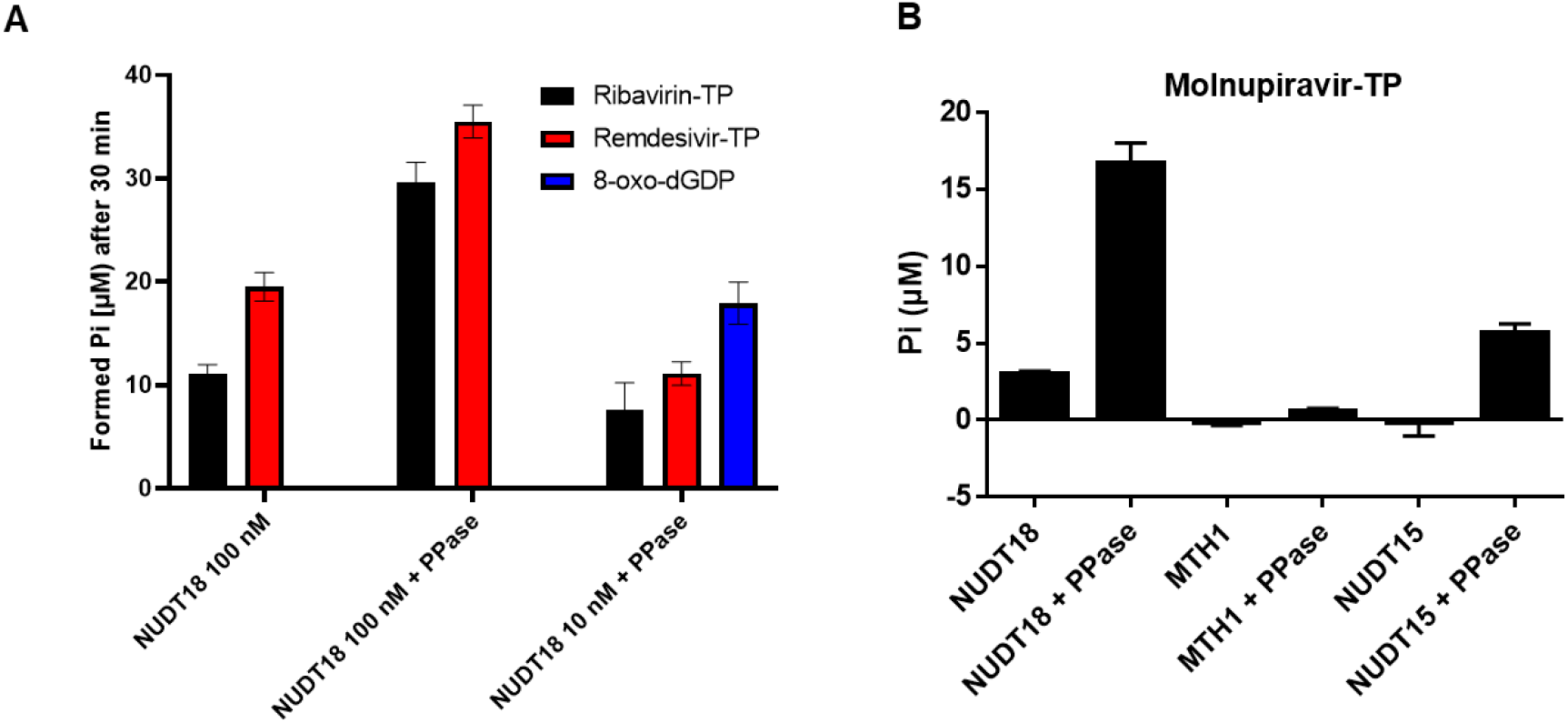
**(A)** Hydrolysis activity of 100 nM NUDT18 with 100 μM Ribavirin-TP or Remdesivir-TP, in presence and absence of 0.2 U/ml *E. coli* PPase, was monitored in assay buffer (20 mM Tris-Acetate pH 8.0, 40 mM NaCl, 10 mM MgAc, 1 mM DTT, 0.05% Tween20) by incubation at 22 °C for 30 minutes. Hydrolysis activity of 10 nM NUDT18 was assessed with 100 μM Ribavirin-TP, Remdesivir-TP or 8-oxo-dGDP in assay buffer with *E. coli* PPase (0.2 U/ml) by incubating reactions at 22 °C for 30 minutes. **(B)** Hydrolysis activity of 20 nM NUDT18, MTH1 and NUDT15 was tested with 100 μM Molnupiravir-TP, in presence and absence of 0.2 U/ml *E. coli* PPase by incubating reactions at 22 °C for 30 minutes. Formed Pi was detected by addition of malachite green reagent and measurement of absorbance at 630 nm.

**Figure 4.**
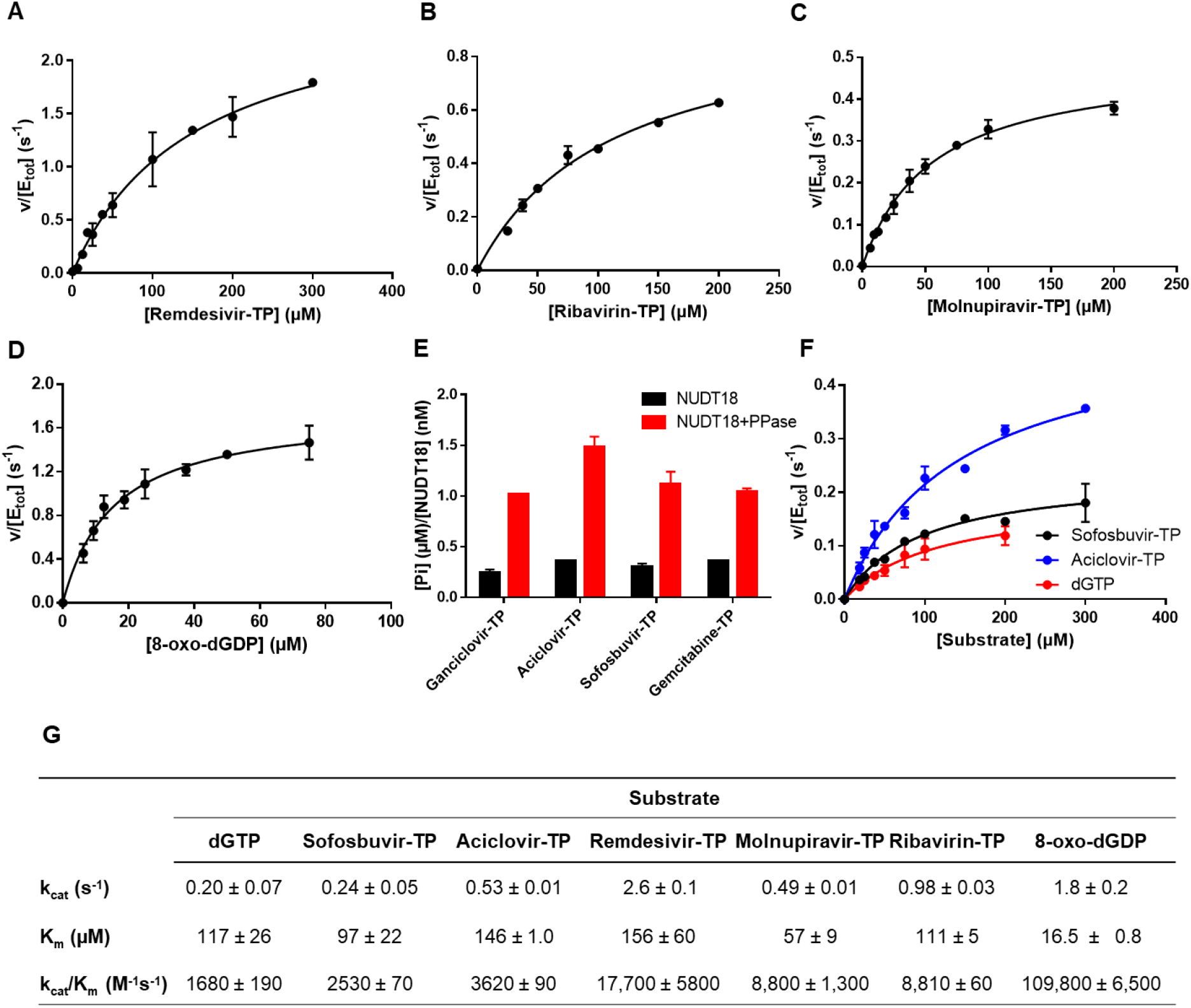
Enzyme saturation curves of NUDT18 with **(A)** Remdesivir-TP, **(B)** Ribavirin-TP, **(C)** Molnupiravir-TP and **(D)** 8-oxo-dGDP. **(E)** Specific activity of 20 nM NUDT18 with 100 μM of triphopshates of the antivirals Ganciclovir, Aciclovir, Sofosbuvir and the cytostatic drug Gemcitabine was tested by incubation for 30 min at 22 °C in assay buffer (20 mM TrisAcetate pH 8.0, 40 mM NaCI, 10 mM MgAcetate, 1 mM DTT and 0.01% Tween 20), in absence and presence of E. coli PPase (0.2 U/ml). Formed Pi was detected using addition of a malachite green reagent followed by measurement of absorbance at 630 nm. Figure shows formed [Pi] (μM) per [NUDT18] (nM). **(F)** Enzyme saturation curves of NUDT18 with Sofosbuvir-TP, Aciclovir-TP and dGTP. Graphs with saturation curves show average of initial rates from two independent experiments with data points performed in duplicate. **(G)** Kinetic parameters of NUDT18 with Remdesivir-TP, Ribavirin-TP, 8-oxodGDP, Aciclovir-TP, Sofosbuvir-TP and dGTP. Presented is average and standard deviation of kinetic parameters determined from two independent experiments and with data points performed in duplicate.

When comparing Remdesivir-TP, Ribavirin-TP and Molnupiravir-TP as substrates for NUDT18, Remdesivir-TP appears to be the preferred substrate. NUDT18 displays a k_cat_ value for Ribavirin-TP hydrolysis that is approximately 1 s^-1^ and a K_m_ value of 111 μM, resulting in a catalytic efficiency value of 8,800 s^-1^M^-1^, which is approximately half of the corresponding value for Remdesivir-TP (Figure 4G). The catalytic efficiency of NUDT18 for Molnupiravir-TP was determined to 8,800 s^-1^M^-1^, which is very close to the corresponding value for Ribavirin-TP. However, the affinity for Molnupiravir-TP seems to be 2-fold higher (K_m_=57 μM) while the turnover number is 2-fold lower (k_cat_=0.49 s^-1^). Comparison of the kinetic parameters of Remdesivir-TP and Ribavirin-TP with those for 8-oxo-dGDP show that the difference primarily lies in the K_m_ value suggesting a better fit of 8-oxo-dGDP in the active site of NUDT18.

### NUDT18 catalyzes the hydrolysis of other antiviral triphosphates

In order to investigate if NUDT18 can catalyze the hydrolysis of other antiviral triphosphates we expanded the substrate screen of NUDT18 and found that Sofosbuvir-TP, Aciclovir-TP, Ganciclovir-TP as well as Gemcitabin-TP are hydrolyzed by NUDT18 to some extent. Similar to the NUDT18-catalyzed hydrolysis of Remdesivir-TP and Ribavirin-TP, hydrolysis takes place at both the β-γ and the α-β phosphate bonds in these substrates with approximately 50% of the hydrolysis taking place at each of these positions (Figure 4E). However, kinetic analysis revealed that the activity with Aciclovir-TP, the best substrate in this panel, is much lower compared to the activity with Remdesivir-TP with a k_cat_/K_m_ value approximately 5 times lower (Figure 4F and G). However, the k_cat_/K_m_ value was 2.2 times higher than the corresponding k_cat_/K_m_ value for dGTP. The fact that Aciclovir contains a guanine moiety like dGTP indicates that the higher activity of NUDT18 with Aciclovir-TP compared to dGTP lies in NUDT18 being able to better accommodate the nucleobase and the modified sugar part of Aciclovir-TP in the active site. This in turn enables a more productive positioning of the phosphate groups for hydrolysis. This is reflected by a 2.6 times higher turnover number (k_cat_) for Aciclovir-TP compared to dGTP, while the K_m_ values are comparable (Figure 4F). However, whether the activity of NUDT18 with these antiviral triphosphates bears clinical relevance for modulating the potency of these antivirals needs further investigation.

### Investigation of binding mode in NUDT18

Remdesivir-TP contains a nitrile group on the C1 atom of the ribose unit (Figure 5). The fact that none of the other tested NUDIX proteins displays any activity with Remdesivir-TP suggests that only the active site of NUDT18 is spacious enough and contains amino acids enabling accommodation of this moiety while still orienting the rest of the molecule allowing for efficient hydrolysis. To date two crystal structures of human NUDT18 have been reported (PDB IDs: 3gg6 and 4hvy), both comprising apo structures of truncated constructs covering residue positions 26-179. Analysis of these structures using SiteMap [SiteMap, Schrödinger, LLC, New York, NY, 2021] indicated that the largest pocket adjacent to the helix containing the NUDIX motif is relatively small and polar in comparison to other NUDIX family members [32]. Docking of Remdesivir-TP to this site did not yield any good-scoring poses with plausible accommodation of the heterocyclic core of Remdesivir-TP. In comparison, the homologous NUDIX family members NUDT1 and NUDT15 contain deeper druggable pockets which readily accommodate nucleobases and small-molecule inhibitors [33, 34]. This suggests that considerable conformational changes are required for NUDT18 to harbor the heterocyclic cores of Remdesivir-TP and other substrates. Attempts to induce a deeper hydrophobic pocket using an induced-fit procedure as described by Loving *et al*. [35] also failed to generate a pocket for Remdesivir-TP with significantly improved SiteScore.

**Figure 5.**
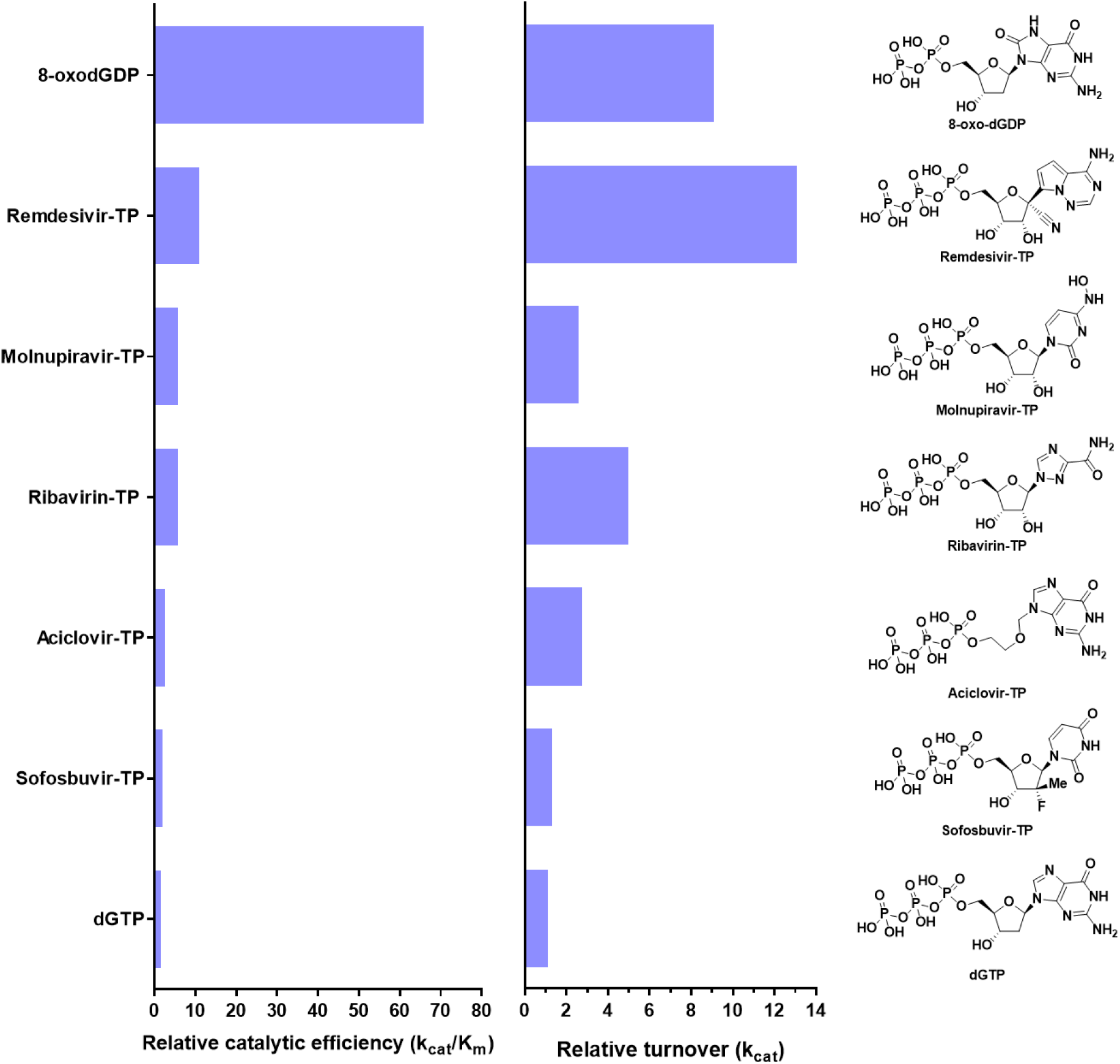
Relative catalytic efficiency and turnover numbers of NUDT18 with tested substrates and their chemical structures.

In order to obtain structural information to understand the binding mode of NUDT18 substrates we carried out extensive crystallization trials of full length NUDT18 without added ligands or together with 8-oxo-dGDP, Remdesivir-TP or Ribavirin-TP. This included seven different commercially available crystallization screens (JCSG+, INDEX, Morpheus, Morpheus II, Pact Premier, Structure Screen and Salt-RX) using different protein concentrations (ranging from 5 mg/mL to 25 mg/mL) and temperatures (4 °C and 20 °C). Unfortunately, we were not able to obtain crystals in any of the conditions tested. As only structures of the nucleosidase binding domain are available for NUDT18 it is possible that the second domain and linking region renders the full-length protein too flexible for crystallization. We performed a structural similarity search using the DALI web server [36], which indicated that hNUDT18 is most structurally similar to human and other eukaryotic MTH1 proteins from dogs, mice and zebrafish (with rms differences of 2.1 to 2.3 Å). As Remdesivir-TP is an analogue of ATP we were particularly interested in comparing the existing hNUDT18 nucleotidase domain structure (PDB ID: 3gg6) with structures of hMTH1 bound to N6-methyl-dAMP (PDB ID: 6qvo, [25] and 2-OH-dATP (PDB ID: 5ghq, [37]. Comparison of these structures showed that residues from the highly conserved NUDIX motif (GX5EX7REVXEEXGU) that coordinate the essential magnesium ions required for substrate hydrolysis superimpose nicely between the structures (Supplementary Figure 1A). This provides some level of certainty as to where the α-phosphate group of Remdesivir-TP would be positioned in hNUDT18 and therefore the orientation of the ligand within the binding pocket. Analysis of the hNUDT18 structure reveals a shallow binding pocket (Supplementary Figure 1B) and structural rearrangements are clearly required to accommodate the adenine base. Analysis of N6-methyl-dAMP and 2-OH-dATP binding in MTH1 shows that the residue W117* forms an important pi-stacking interaction with the adenine base. In NUDT18 the corresponding residue is an alanine, however a tyrosine residue (Y72, N33* in hMTH1) which would be capable of performing an equivalent pi-stacking interaction is situated in close proximity (Supplementary Figure 1C). In the NUDT18 apo structure Y72 clashes with the bound base when compared to the positioning of the ligands in the hMTH1 structures. A conformational adjustment must occur in order to make room for the substrate. Repositioning of this tyrosine would shape the binding pocket in such a way that it, while still being shallow, is capable of accommodating N6-methyl-dAMP and 2-OH-dATP and importantly, Remdesivir-TP with its additional nitrile group (Supplementary Figure 1D). The lower K_m_ value of 8-oxo-dGDP for NUDT18 suggests it fits better within the active site of the enzyme than Remdesivir-TP. In the absence of a full-length NUDT18 structure, a structure of the NUDT18 nucleotidase domain bound to the natural substrate 8-oxo-dGDP or to Remdesivir-TP would be very useful for elucidating the exact structural rearrangements that occur within the nucleotide binding site upon binding of these ligands.

In an attempt to understand which structural features of the substrates that influence their quality as NUDT18 substrates, we compared their values of catalytic efficiency and turnover numbers of NUDT18 catalyzed hydrolysis for the tested substrates relative to the corresponding values for dGTP and examined the chemical structures of the tested substrates (Figure 5). NUDT18 has previously been reported to catalyze the hydrolysis of 8-oxo-dGDP and 8-oxo-GDP with the same efficiency [28], demonstrating that the 2’-OH group can be accommodated in the active site of NUDT18. This suggests a flexibility towards alterations within this part of the nucleoside substrate. Among the tested substrates, the turnover number is highest for Remdesivir-TP suggesting that either NUDT18 catalyzed hydrolysis or product release is faster for Remdesivir-TP compared to 8-oxo-dGDP under substrate saturating conditions. However, the K_m_ value is considerably lower for 8-oxo-dGDP (Figure 4G), indicating specific interactions between the oxidized nucleobase and active site residues of NUDT18. The lower K_m_ value, which translates into higher catalytic efficiency for 8-oxo-dGDP, may in part be due to nucleoside triphosphates binding with lower affinity to the active site compared to nucleoside diphosphates. The difference in catalytic efficiency between Remdesivir-TP and Ribavirin-TP can likely be attributed to differences in positioning for efficient hydrolysis and/or product release since their K_m_ values are similar. The lower k_cat_ value for Molnupiravir-TP compared to Remdesivir-TP indicates a less efficient binding of Molnupiravir-TP for productive catalysis. The NUDT18-catalyzed hydrolysis of Molnupiravir-TP seems to take place mostly at the α-β phosphate bond while in the case of Remdesivir-TP hydrolysis occurs at both the β-γ and the α-β phosphate bonds to approximately the same extent (Figure 3A and B) suggesting that accommodation of the N4-hydroxycytidine moiety of Molnupiravir-TP in the active site is more restricted in comparison to binding of the bicyclic triazin moiety of Remdesivir-TP. Interactions between the nucleobase and/or the nitrile group of Remdesivir-TP and NUDT18 may contribute to a more productive binding. However, in absence of structural information, it is difficult to draw accurate conclusions about the binding mode. Nevertheless, the fact that the active site of NUDT18 can accommodate molecules with rather different structures and that hydrolysis takes place at two positions with equal efficiency, with the exception of Molnupiravir-TP, indicates a relatively flexible binding mode for these substrates.

### Inhibition of NUDIX enzymes by Remdesivir and metabolites

Since Remdesivir-TP is a substrate for NUDT18 we argued that Remdesivir and its metabolites may inhibit NUDT18 and related NUDIX proteins. We therefore tested MTH1 and NUDT15 for inhibition by Remdesivir, the Remdesivir metabolite GS-441524 (arising from cellular hydrolysis of Remdesivir) and Remdesivir-TP. We tested NUDT18 for inhibition by Remdesivir and GS-441524. NUDT15 was found to be modestly inhibited by 100 μM Remdesivir (26%) as well as by GS-441524 (21 %) (Figure 6). MTH1 was slightly inhibited by GS-441524 (23%) while no inhibition of NUDT18 by any of the tested Remdesivir derivatives was observed. The modest inhibitory effect of Remdesivir and its metabolites imply that Remdesivir treatment is unlikely to have a pronounced effect on the cellular activity of these enzymes.

**Figure 6.**
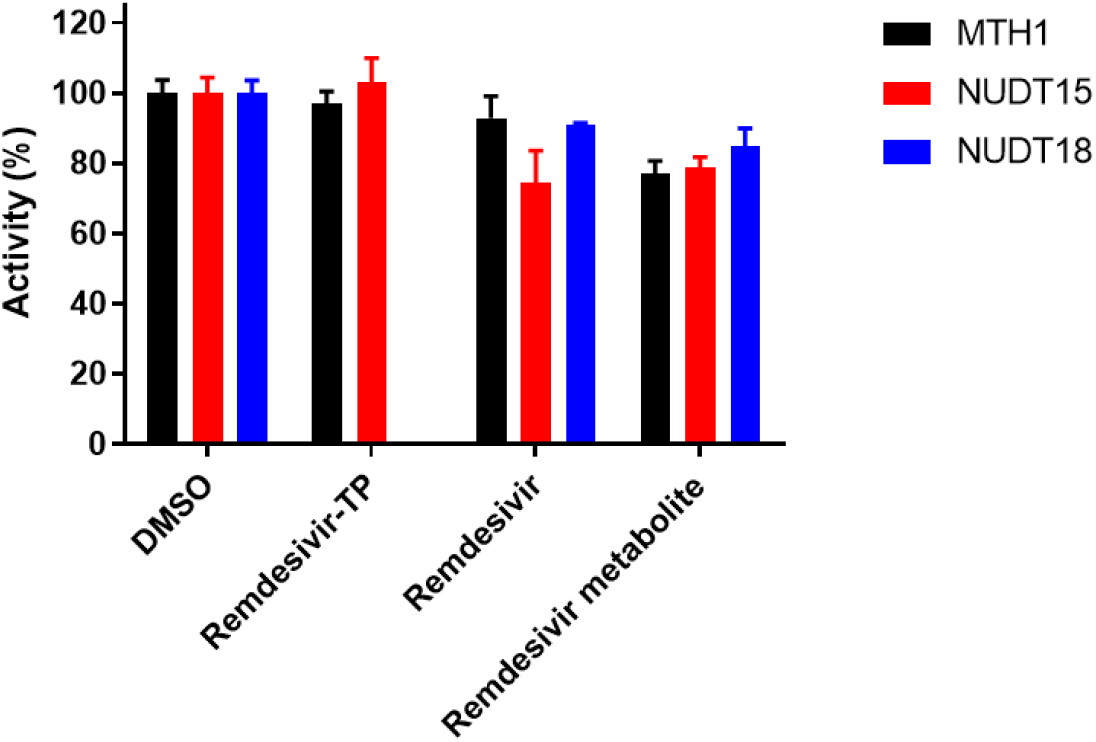
Inhibition of NUDIX enzymes by Remdesivir and its metabolites. MTH1 and NUDT15 was tested for inhibition by 100 μM Remdesivir-TP, Remdesivir or Remdesivir metabolite (GS-441524) in an activity assay using 100 μM dGTP as substrate and 10 nM NUDT15 or MTH1. The potency of Remdesivir and Remdesivir metabolite (GS-441524) to inhibit NUDT18 was tested using μM 8-oxo-dGDP as substrate and using 100 nM NUDT18. NUDT15 is modestly inhibited by Remdesivir and activity of all three enzymes are reduced by approximately 20% in presence of the Remdesivir metabolite (GS-441524).

### Consequence of NUDT18 activity with antiviral triphosphates

The results presented here suggest that NUDT18 can convert Remdesivir-TP to Remdesivir-DP and Remdesivir-MP in the cell. However, since Remdesivir-DP and Remdesivir-MP are substrates for adenylate kinase 2 they will be phosphorylated and a cellular equilibrium between production of Remdesivir-TP by kinases and hydrolysis catalyzed by NUDT18 is likely established. Furthermore, the action of intracellular 5’-nucleotidase will affect the concentration of Remdesivir-MP. In a similar way the cellular concentrations of the metabolites of Molnupiravir and Ribavirin will be regulated by the activities of adenylate kinase 2 and NUDT18 and in the case of Ribavirin also ITPase. For NUDT18 to play a clinically relevant role in modulating Remdesivir efficacy it has to be present in the cells that are infected by the virus. Importantly, NUDT18 is ubiquitously expressed, displays low tissue specificity (https://www.proteinatlas.org/ENSG00000275074-NUDT18) and is expressed in lung and in respiratory epithelial cells (https://www.ebi.ac.uk/gxa/home). This suggests that NUDT18 would be able to modulate the effective concentration of Remdesivir-TP and Molnupiravir-TP at their intended site of action. Potentially, inhibition of NUDT18 would be a means to increase the efficacy of these drugs. However, no NUDT18 inhibitors enabling such investigations are available to the best of our knowledge.

The importance of intracellular metabolism for efficiency of antivirals is exemplified by the activity of human ITPase with Ribavirin-TP [22]. Approximately a third of the human population carries low activity ITPase alleles [22] and it is clear that cellular ITPase activity influences the antiviral efficacy of Ribavirin as well as its adverse effects. Similarly, higher expression of NUDT18 in respiratory epithelial cells in a COVID-19 patient, potentially resulting in considerable Remdesivir-TP and Molnupiravir-TP hydrolysis, may result in lower efficiency of the corresponding antivirals compared to in a patient with lower NUDT18 expression levels. This may explain differences in response to Remdesivir treatment between patients.

Here we show that human NUDT18 can catalyze the hydrolysis of the active metabolites of the antivirals Remdesivir, Ribavirin and Molnupiravir. These novel NUDT18 activities may be clinically relevant and warrants further investigation and emphasizes the need for NUDT18-specific inhibitors enabling studies of the cellular NUDT18 function and role in modulating the efficacy of these antiviral drugs.

In summary, we suggest a cellular role of NUDT18 as a sanitizer of modified nucleotides, including antiviral triphosphates.

## Materials and methods

### Protein production

MTH1, NUDT15, NUDT18, NUDT5 and NUDT9, and *E. coli* PPase were produced as N-terminally His-tagged proteins and were expressed and purified by the Protein Science Facility, Karolinska Institute, using Ni-IMAC (HisTrap HP, GE Healthcare) followed by gel filtration (HiLoad 16/60 Superdex 75 or HiLoad 16/60 Superdex 100, GE Healthcare) as previously described [38]. Proteins were concentrated and then stored as aliquots at −80 °C in storage buffer (20 mM HEPES pH 7.5, 300 mM NaCl, 10% glycerol and 0.5 mM TCEP (tris-(carboxyethyl)phosphine)). ITPase, dUTPase and dCTPase were produced as N-terminally His-tagged proteins and expressed and purified as earlier described [33].

### Screen of human hydrolases for Remdesivir-TP activity

Hydrolysis activity of MTH1, NUDT15, NUDT18, NUDT5 and NUDT9 (200 nM) with 100 μM Remdesivir-TP was assayed in reaction buffer (20 mM Tris-Acetate pH 8.0, 40 mM NaCl, 10 mM MgAc, 1 mM DTT, 0.01% Tween20) by incubating reaction mix in wells of clear 96-well plates (Nunc 269620) at 22°C for 30 minutes. Formed Pi was detected by addition of malachite green reagent [39] and measurement of absorbance at 630 nm. Activity of 200 nM MTH1, NUDT15, NUDT18 and NUDT5, with 100 μM Remdesivir-TP was also tested in the presence of *E. coli* PPase (0.2 U/ml) which converts formed PPi to Pi. Activity of the human pyrophoshates: ITPase, dUTPase and dCTPase with 100 μM Remdesivir-TP in the presence of PPase (0.2 U/ml) was assayed and compared to activity of NUDT18 with 100 μM 8-oxo-dGDP. Buffer controls with and without PPase were included and the background signal was subtracted from the assay data.

### Assessment of specific activity of NUDT18 with Remdesivir-TP, Ribavirin-TP and 8-oxo-dGDP

In order to test if NUDT18 can also hydrolyze Ribavirin-TP and to compare its activity towards Remdesivir-TP (BioCarbosynth) and 8-oxo-dGDP (Jena Bioscience), 100 μM Ribavirin-TP (Jena Bioscience), Remdesivir-TP or 8-oxo-dGDP, respectively, was incubated with 10 nM NUDT18 in reaction buffer (20 mM TrisAcetate pH 8.0, 40 mM NaCl, 10 mM MgAc, 1 mM DTT, 0.01% Tween20) fortified with 0.2 U/ml PPase for 30 minutes at 22 °C after which formed Pi was detected by addition of malachite green reagent and measurement of absorbance at 630 nm in a Hidex spectrophotometer.

### Assessment of specific activity with Molnupiravir-TP

In order to test the activity of NUDT18 with the active metabolite of the orally active antiviral Molnupiravir, Molnupiravir-TP, and to investigate if the closely related NUDIX enzymes NUDT15 and MTH1 can hydrolyze this substrate hydrolysis activity of 20 nM NUDT18, MTH1 and NUDT15 was tested with 100 μM Molnupiravir-TP, in the presence and absence of 0.2 U/ml *E. coli* PPase. Reactions were incubated at 22 °C for 30 minutes and formed Pi was detected by addition of malachite green reagent and measurement of absorbance at 630 nm.

### Determination of kinetic parameters of NUDT18 with Remdesivir-TP, Ribavirin-TP and Molnupiravir-TP

Initial rates of NUDT18 (10 nM) were determined in assay buffer (20 mM TrisAcetate pH 8.0, 40 mM NaCl, 10 mM MgAc, 1 mM DTT, 0.01% Tween20, 0.2 U/ml PPase) at concentrations ranging from 0 to 300 μM for Remdesivir-TP and from 0 to 200 μM for Ribavirin-TP and Molnupiravir-TP. Reaction mixtures were incubated for 0, 10, 20 and 30 minutes. Formed Pi was detected by adding malachite green reagent and measurement of absorbance at 630 nm. A Pi standard curve was included on the plate and used to convert absorbance to Pi concentration. Enzyme saturation curves were also produced with 8-oxo-dGDP, which has been described to be the endogenous substrate of NUDT18, using the same enzyme concentration and buffer conditions. 8-oxo-dGDP concentrations ranging between 0 and 100 μM were used. Initial rates were calculated using linear regression and plotted against substrate concentration. The Michaelis-Menten equation was fitted to the data points and kinetic parameters of NUDT18 were determined using GraphPad Prism 8.0. Data points were measured in duplicate and two independent experiments were performed.

### Assessment of specific activity of NUDT18 with triphosphates of other antivirals

The finding that NUDT18 displays pronounced activity with the antiviral triphosphates Remdesivir-TP, Ribavirin-TP and Molnupiravir-TP prompted us to also test the activity of NUDT18 with an expanded panel of antiviral triphosphates that are considered to be the active antiviral metabolite. The panel included the triphosphates of; Sofosbuvir used for the treatment of hepatitis C (Sofosbuvir-TP, (2R)-(2’-deoxy-2’-fluoro-2-methyl-uridine-5’-TP, Sierra bioresearch), Ganciclovir used to treat cytomegalovirus infection treatment (Ganciclovir-TP, Sierra bioresearch) and Aciclovir used for Herpes simplex virus infection therapy, (Aciclovir-TP, Sierra bioresearch). We also included the active metabolite of the cytostatic drug gemcitabine (Gemcitabine-TP, Sierra bioresearch). NUDT18 (20 nM) was incubated with 100 μM of Ganciclovir-TP, Aciclovir-TP, Sofosbuvir-TP and Gemcitabine-TP for 30 min at 22 °C in assay buffer (20 mM TrisAcetate pH 8.0, 40 mM NaCl, 10 mM Magnesium Acetate, 1 mM DTT and 0.01% Tween 20), in presence and absence of PPase 0.2 U/ml. Formed Pi was detected by addition of malachite green reagent followed by measurement of absorbance at 630 nm.

### Determination of kinetic parameters of NUDT18 with triphosphates of antiviral compounds

Initial rates of NUDT18 (10 nM) were determined in assay buffer (20 mM Tris-Acetate pH 8.0, 40 mM NaCl, 10 mM MgAc, 1 mM DTT, 0.01% Tween20, 0.2 U/ml PPase at concentrations ranging from 0 to 300 μM for Sofosbuvir-TP and Aciclovir-TP by incubating the reactions for 0, 10, 20 and 30 minutes. Formed Pi was detected by adding malachite green reagent and measurement of absorbance at 630 nm. The concentration of formed Pi was calculated using a Pi standard curve included on the assay plate. Enzyme saturation curves were also produced for dGTP (GE Healthcare #27-1870-04) using a concentration range between 0 and 200 μM. Initial rates were plotted against substrate concentration and the Michaelis-Menten equation was fitted to the data points allowing kinetic parameters of NUDT18 to be determined. Data points were measured in duplicate and two independent experiments were performed.

### Computational analysis

All calculations were performed using Schrödinger Suite 2021-1 [Schrödinger LLC, New York, NY, 2021].

#### Protein structure preparation

The apo structure of human NUDT18 (3GG6.pdb, unpublished) was prepared using the Protein Preparation Wizard. Briefly, the raw PDB structure was processed by automatically assigning bond orders, adding hydrogens, converting seleno-methionine to methionine, adding missing side chains, and creating possible disulfide bridges. Residues with alternate positions were locked in the conformations with the highest average occupancy. Hydrogen bonding networks were optimized automatically by optimization of hydroxyls, Asn, Gln, and His residue states using ProtAssign. A restrained minimization was then performed using the OPLS4 force field, until an RMSD convergence of 0.30 Å was reached for the heavy atoms.

#### Binding site analysis

SiteMap was used to probe the prepared NUDT18 structure for possible ligand binding sites. The 5 top-ranked potential binding sites were identified. At least 15 site points per reported sites were required. The more restricted definition of hydrophobicity together with a standard grid (0.7 Å) were used. Site maps at 4 Å or more from the nearest site points were cropped.

#### Ligand structure preparation

The structure of Remdesivir-TP was copied as isomeric smiles from PubChem (CID: 56832906) and pasted as 3D structure into Maestro. Ligprep was then used to prepare the structure of Remdesivir-TP for docking. The OPLS4 force field was used for minimizations; possible ionization states at pH 7.0 ± 2.0 were generated using Epik, and possible tautomers were generated; specified chiralities were retained and at most 1 stereoisomer was generated.

#### Induced-fit docking

Remdesivir-TP was docked to NUDT18 using the Induced-Fit Docking protocol [40]. The site points of largest site identified by SiteMap were used to center the docking boxes. The size of the enclosing box was set to 20 Å. Glide XP was used for the redocking step, otherwise all settings were kept at default values.

### Inhibition assay

Since Remdesivir-TP acts as a substrate of NUDT18 we decided to investigate if Remdesivir or its metabolites act as inhibitors of NUDT18 or its close relatives NUDT15 and MTH1. Activity of MTH1 (10 nM) and NUDT15 (10 nM) was tested in assay buffer (0.1 M Tris-Acetate pH 7.5, 40 mM NaCl, 1 mM DTT, 10 mM Mg-Acetate) fortified with PPase (0.2 U/ml), with 100 μM dGTP in the presence of 1% DMSO or with 100 μM of Remdesivir-TP (#G167050, BioCarbosynth), Remdesivir (#AG170167, BioCarbosynth) or Remdesivir metabolite (#AG167808, BioCarbosynth). NUDT18 activity was assayed using 100 nM NUDT18 and 100 μM 8-oxo-dGDP in assay buffer. The reaction mixture was incubated for 30 min at 22°C after which formed Pi was detected by addition of malachite green reagent followed by absorbance measurements at 630 nm in a Hidex spectrophotometer.

## Acknowledgements

The authors thank the Protein Science Facility at Karolinska Institutet/SciLifeLab (http://ki.se/psf) for protein production and Mari Kullman Magnusson for excellent laboratory support and Louise Sjöholm and Athina Pliakou for administrative support. We thank Dr Maeve Long for linguistic corrections. This work was supported by the European Research Council (TAROX Programme, ERC-695376 to TH), the Swedish Research Council (2015-00162, 2017-06095 to TH and 2018-03406 to P.S.), the Torsten and Ragnar Söderberg Foundation (TH), the Knut and Alice Wallenberg Foundation (KAW2014.0273 to T.H.), the Swedish Cancer Society (CAN2018/0658 to T.H. and 20 1287 PjF to P.S.), KI funds (2020-02211 to A.S.J), the Alfred Österlund foundation (to P.S), the Åke Olsson foundation for haematological research (2020-00306 to M.M.), and the EU/EFPIA/OICR/McGill/KTH/Diamond Innovative Medicines Initiative 2 Joint Undertaking (EUbOPEN grant no 875510, M.M. and E.J.H.). Funding for open access charge: The Swedish Research Council.

## Author contributions

A.S.J wrote the manuscript with input from E.S., E.H. and M.M. All authors commented on the manuscript. E.H., M.M. and A.-S.J. conceived the project. A.S.J. designed, performed and analysed all biochemical experiments. E.J.H. and E.S. designed, performed and analysed computational chemistry experiments and structural analyses. P.S., T.H. and M.M. supervised the project.

## Data availability

All data are available from the authors upon request.

---

**Supplementary Figure 1.**
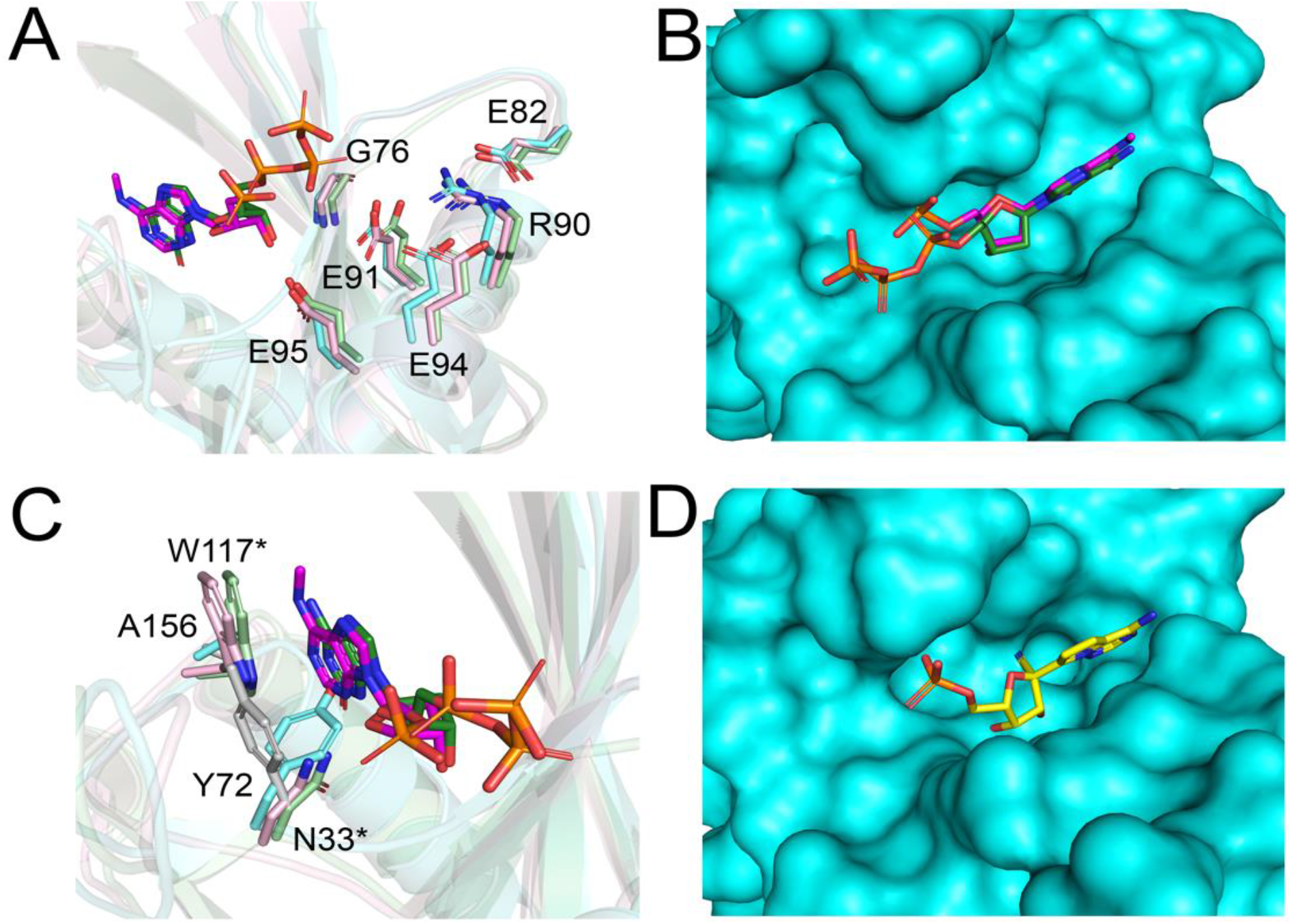
Structural comparison of hNUDT18 with hMTH1. (A) Superposition of the hNUDT18 nucleotidase domain (cyan, residues 36-179, PDB ID: 3gg6) with hMTH1 bound to N6-methyl-dAMP (light pink, PDB ID: 6qvo, 10.1074/jbc.RA120.012636) and hMTH1 bound to 2-OH-dATP (pale green, PDB ID: 5ghq, (31). N6-methyl-dAMP and 2OH-dATP are shown as magenta and dark green stick models, respectively. Residues from the NUDIX box motif from MTH1 and NUDT18 that coordinate magnesium, and therefore facilitate substrate hydrolysis, are shown as sticks. Numbering corresponds to residues in the NUDT18 sequence. (B) Surface representation of NUDT18 (cyan) superimposed with hMTH1-N6-methyl-dAMP and hMTH1-2-OH-dATP, showing NUDT18 has a shallow binding pocket. (C) Binding pocket surrounding the adenine base. A tryptophan residue (W117*) from hMTH1 involved in an important pi-stacking interaction is highlighted. Asterisks represent residues from the hMTH1 sequence. The residue Y72 in hNUDT18 needs to reposition (shown in grey) in order to accommodate the adenine base in the binding pocket. (D) Surface representation showing the nucleotide binding pocket after repositioning of Y72. Remdesvir-MP (yellow stick representation) has been positioned in the binding pocket assuming a similar binding mode to N6-methyl-dAMP and 2-OH-dATP in the hMTH1 structures.

